## Supplemental Material for "Cotyledon opening during seedling deetiolation is determined by ABA-mediated splicing regulation"

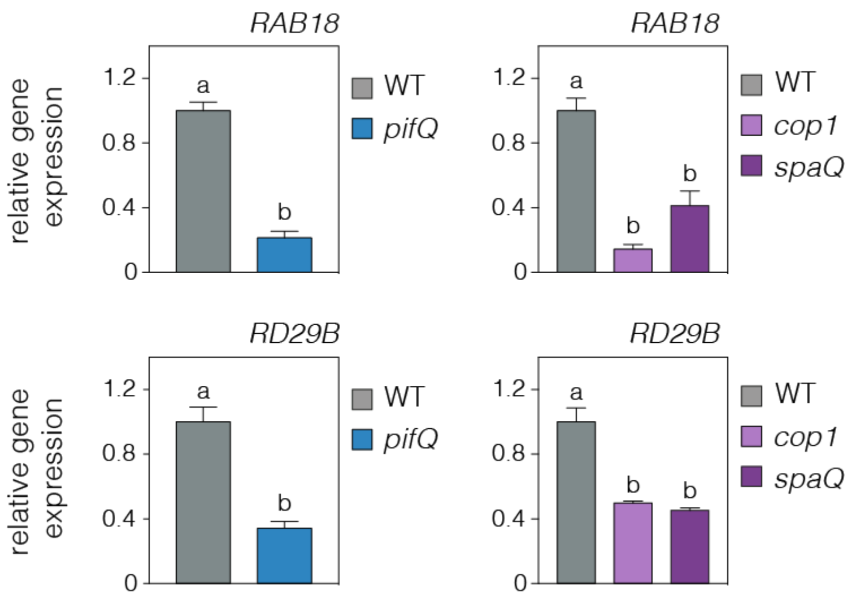


**Appendix Fig. S1. *RAB18* and *RD29B* expression levels in constitutive photomorphogenic mutants.** *RAB18* and *RD29B* transcript levels, quantified from publicly available RNA sequencing data, of *pifQ*, *cop1* and *spaQ* seedlings and their respective wild type (WT) grown for 3 days in the dark. Data are the means ± SE of biological triplicates, with different letters indicating statistically significant differences between genotypes ($P < 0.05$) by unpaired *t* test (left) and Tukey's multiple comparison test (right). RNA sequencing data were obtained from GSE112662 (Pham et al., 2018) and GSE164122 (Martín and Duque, 2021).

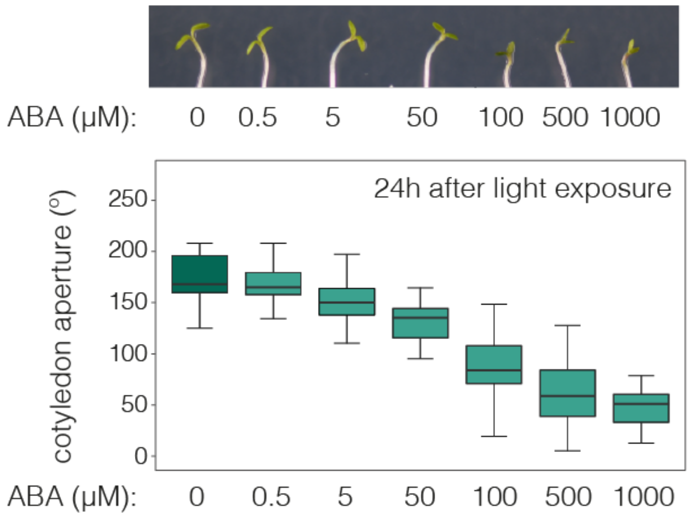


**Appendix Fig. S2. ABA repression of cotyledon aperture in the light.** Representative image (top) and quantification of the cotyledon aperture (bottom) in at least 35 wild-type seedlings grown for 3 days in the dark and then exposed to white light in the presence of different concentrations of ABA. Cotyledon aperture is shown as the difference between the cotyledon angle of each seedling at 24 hours of ABA treatment and the median of the cotyledon angle at time 0 hours. The experiment was repeated at least twice with similar results.

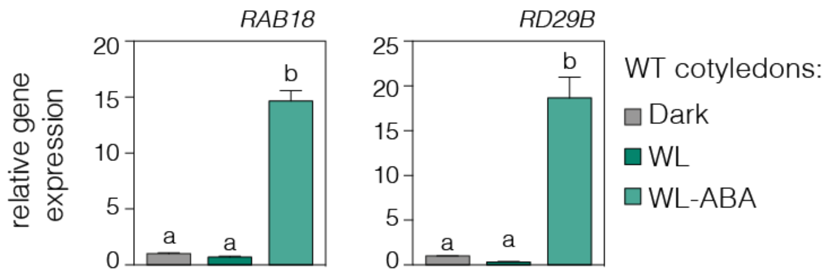


**Appendix Fig. S3. *RAB18* and *RD29B* expression levels in ABA-treated cotyledons during seedling deetiolation.** *RAB18* and *RD29B* transcript levels, quantified from our RNA sequencing data, in wild-type (WT) cotyledons from seedlings grown for 3 days in the dark and then exposed to continuous white light (WL) for 3 hours in the absence or presence of ABA (100 µM). Data are the means ± SE of biological triplicates and relative to the dark timepoint. Different letters indicate statistically significant differences between conditions by Tukey's multiple comparison test ($P < 0.05$).

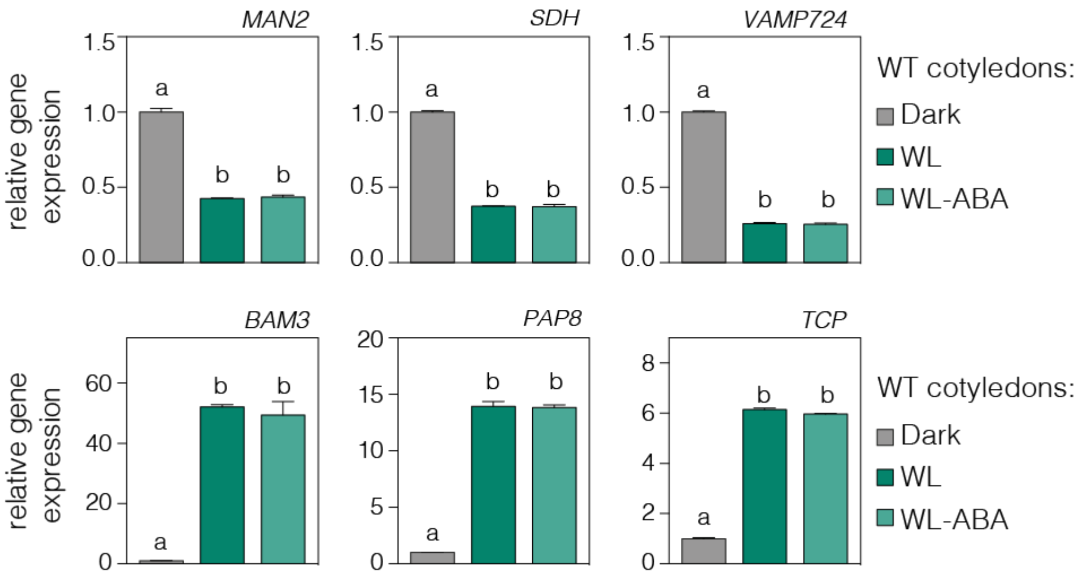


**Appendix Fig. S4. Expression levels of genes whose light responsiveness is unaffected by ABA.** Transcript levels, quantified from our RNA sequencing data, in wild-type (WT) cotyledons from seedlings grown for 3 days in the dark and then exposed to continuous white light (WL) for 3 hours in the absence or presence of ABA (100 µM). Data are the means ± SE of biological triplicates and relative to the dark timepoint. Different letters indicate statistically significant differences between conditions by Tukey's multiple comparison test ($P < 0.05$).

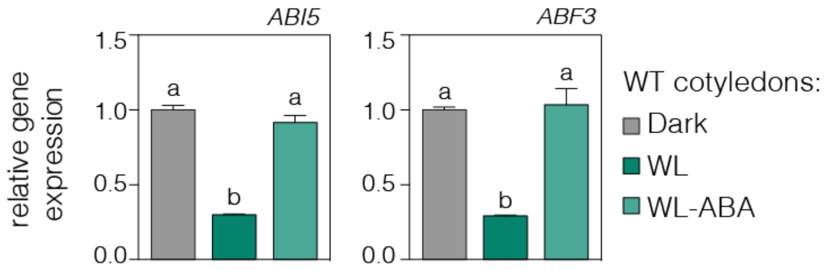


**Appendix Fig. S5. *ABI5* and *ABF3* expression levels in ABA-treated cotyledons during seedling deetiolation.** *ABI5* and *ABF3* transcript levels, quantified from our RNA sequencing data, in wild-type (WT) cotyledons from seedlings grown for 3 days in the dark and then exposed to continuous white light (WL) for 3 hours in the absence or presence of ABA (100 μM). Data are the means ± SE of biological triplicates and relative to the dark timepoint. Different letters indicate statistically significant differences between conditions by Tukey's multiple comparison test ($P < 0.05$).

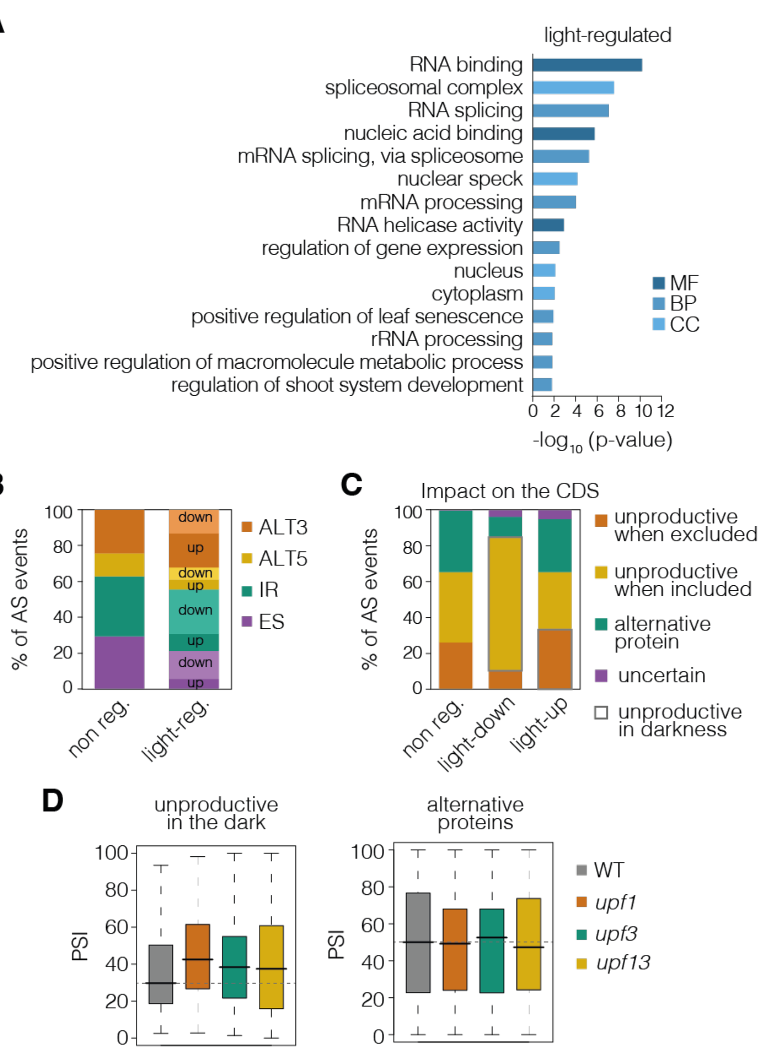


**Appendix Fig. S6. Light-regulated splicing changes in cotyledons from etiolated seedlings. (A)** Top 15 enriched gene ontology categories of molecular function (MF), biological process (BP) and cellular component (CC) for the 224 genes defined as differentially spliced in response to white light in cotyledons. DAVID *p*-value indicates significance (Fisher's exact test; $P < 0.05$; Dataset EV4). **(B)** Number of events by AS type for which the inclusion of the alternative sequence is up- or downregulated by light in comparison to non-regulated AS events (non reg.). ALT5, alternative 5´splice site; ALT3, alternative 3´splice site; IR, intron retention; ES, exon skipping. **(C)** Percentage of AS events located in gene coding sequence (CDS) regions that potentially generate unproductive mRNAs or alternative protein isoforms (see Methods for details) in three groups of AS events: non-regulated (non-reg.), up-, or downregulated by light. AS events that have the ultimate effect of generating unproductive isoforms in the dark are highlighted. **(D)** Percent of inclusion (PSI) values for the CDS-located light-regulated AS events that generate unproductive transcripts in the dark (left) or alternative proteins (right) in wild-type (WT), *upf1*, *upf3* and *upf1upf3* seedling samples. This quantification was conducted with RNA sequencing data obtained from GSE41432 (Drechsel et al., 2013).

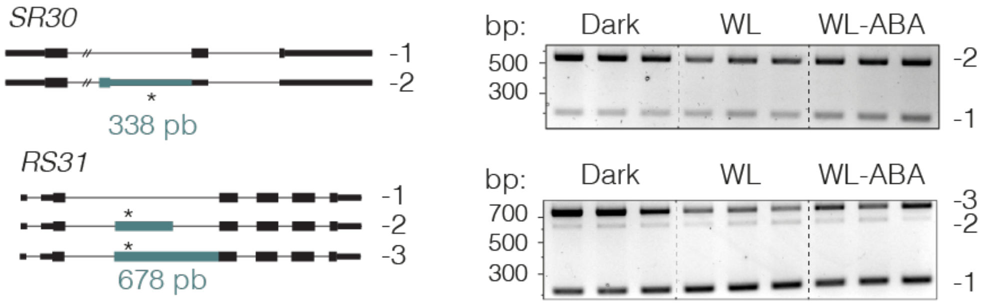


**Appendix Fig. S7. Light and ABA regulation of AS events in *SR30* and *RS31*.** RT-PCR analysis of *SR30* (top) and *RS31* (bottom) alternative transcript levels in seedlings grown for 3 days in the dark and then exposed to continuous white light (WL) for 8 hours (h). The three gel lanes for each condition are biological triplicates, and the gene diagrams on the left show the alternative sequences in green, with asterisks indicating the location of in-frame premature stop codons. bp, base pairs.

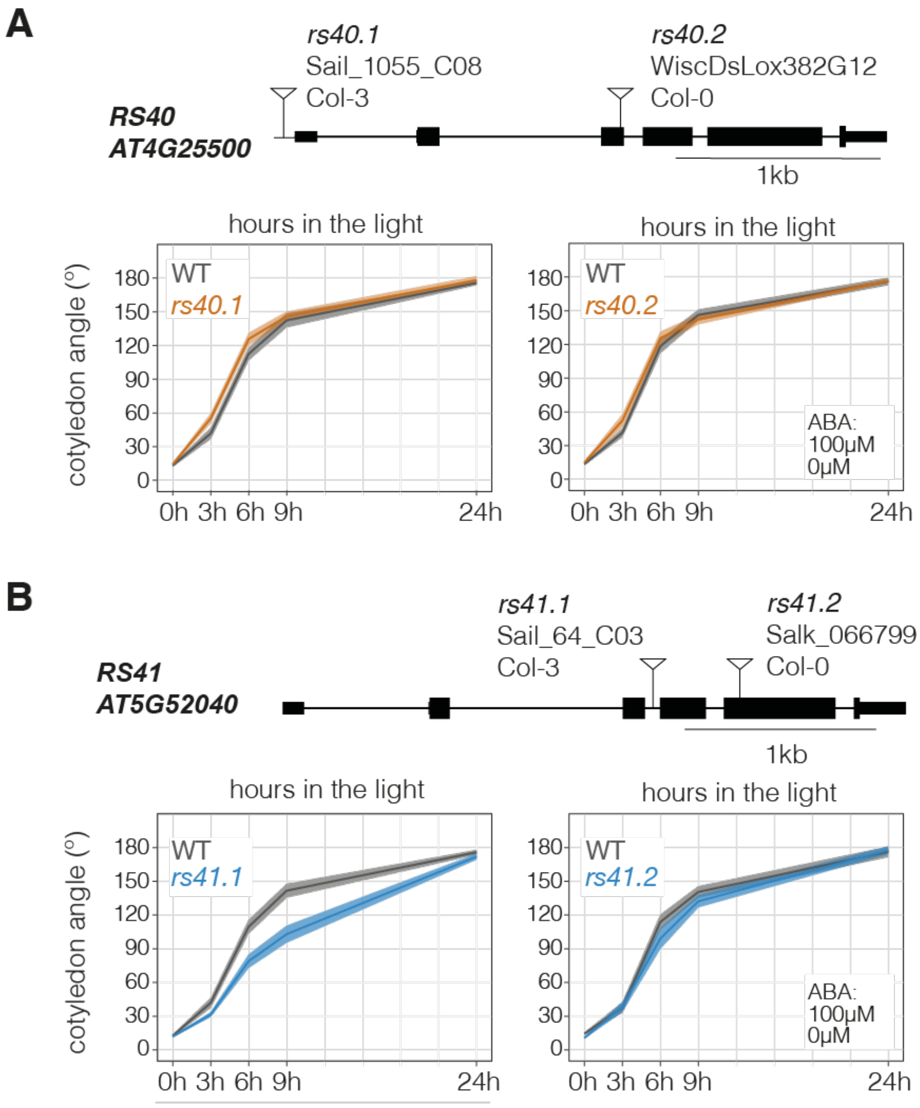


**Appendix Fig. S8. Light regulation of cotyledon opening in *rs40* and *rs41* single mutants.** For *RS40* **(A)** and *RS41* **(B),** gene diagrams (top) indicating the T-DNA insertion sites for two mutant lines per gene and the corresponding cotyledon phenotypes (bottom) are shown. Quantification of cotyledon opening was conducted in each mutant and its respective wild-type (WT) background (Col-0 or Col-3). Seedlings were grown for 3 days in the dark and then exposed to white light for 3, 6, 9 or 24 hours (h). Thick lines and shaded areas represent respectively the median and the interquartile range of at least 45 seedlings. The experiment was repeated at least twice with similar results.

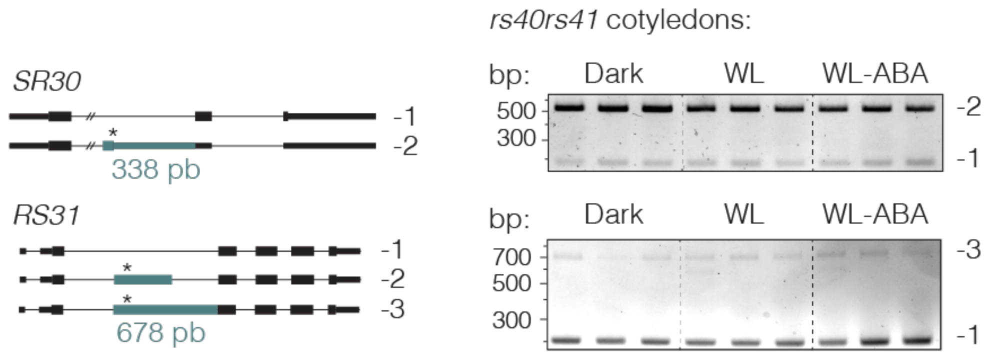


**Appendix Fig. S9. Light and ABA regulation of AS in the *rs40 rs41* double mutant.** RT-PCR analysis of *SR30* (top) and *RS31* (bottom) alternative transcript levels in *rs40rs41.1* cotyledons of seedlings grown for 3 days in the dark and then exposed to continuous white light (WL) for 8 hours in the absence or presence of ABA (100 µM). The three gel lanes for each condition are biological triplicates, and the gene diagrams on the left show the alternative sequences in green, with asterisks indicating the location of in-frame premature stop codons. bp, base pairs.

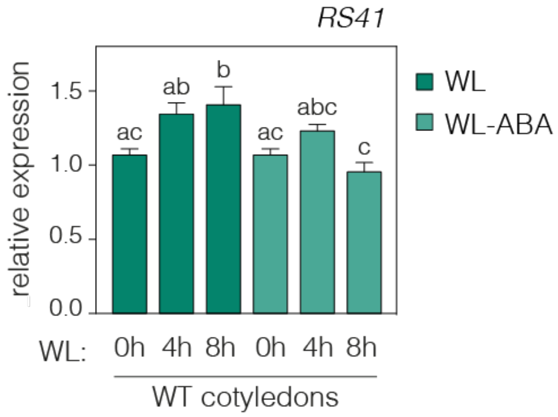


**Appendix Fig. S10. ABA regulation of *RS41* expression during seedling deetiolation.** RT-qPCR analysis of *RS41* transcript levels in cotyledons of wild-type (WT) seedlings grown for 3 days in the dark and then exposed to continuous white light (WL) for 4 or 8 hours (h) in the absence or presence of ABA (100 µM). *PP2A* was used as a reference gene, and expression levels in the dark-grown (0h WL) were set to 1. Data are the means ± SE of biological triplicates, and different letters indicate statistically significant differences between conditions by Tukey's multiple comparison test ($P < 0.05$).

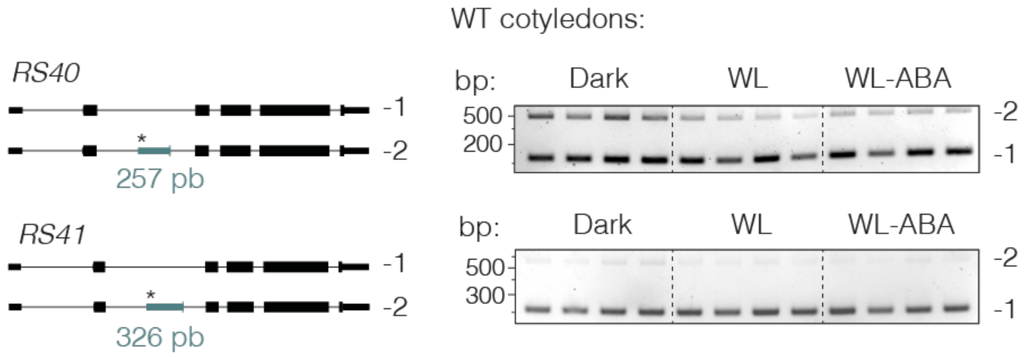


**Appendix Fig. S11. Light and ABA regulation of AS events in *RS40* and *RS41*.** RT-PCR analysis of *RS40* (top) and *RS41* (bottom) alternative transcript levels in wild-type (WT) cotyledons of seedlings grown for 3 days in the dark and then exposed to continuous white light (WL) for 3 hours (h) in the absence or presence of ABA (100 μM). The four gel lanes for each condition are biological quadruplets, and the gene diagrams on the left show the alternative sequences in green, with asterisks indicating the location of in-frame premature stop codons. bp, base pairs.

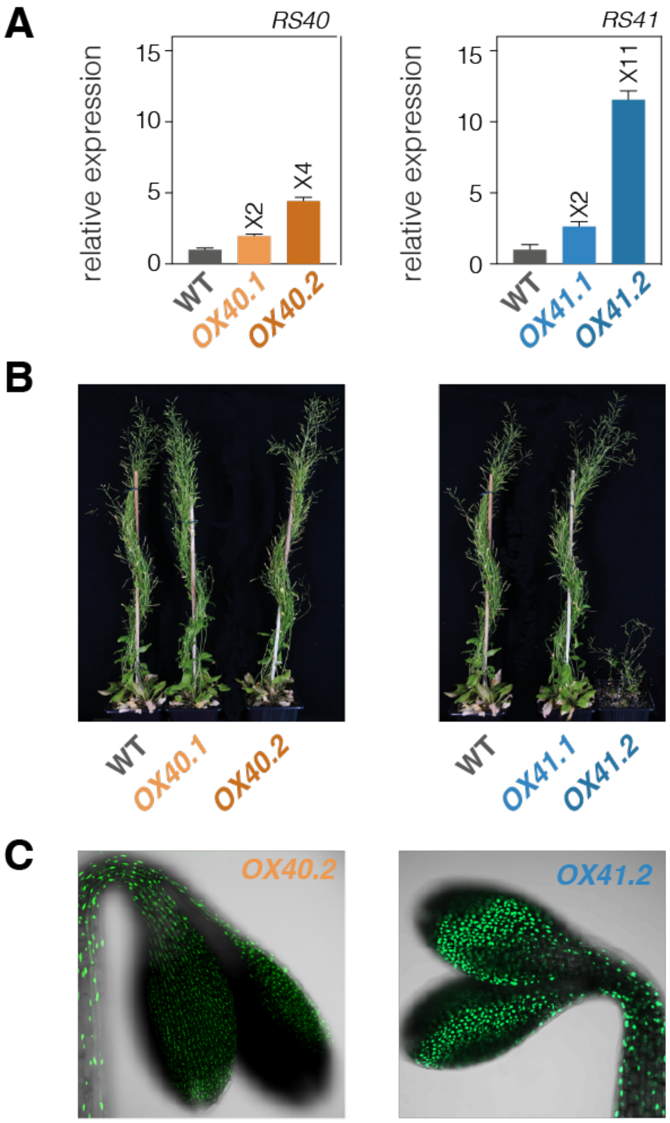


**Appendix Fig. S12. Generation of transgenic plants overexpressing *RS40* and *RS41*. (A)** RT-qPCR analysis of *RS40* and *RS41* transcript levels in the wild-type (WT) and two transgenic overexpression lines grown for 3 days in the dark and then exposed to continuous white light for 6 hours (h). *PP2A* was used as a reference gene, and expression levels in the WT were set to 1. Data are the means ± SE of technical triplicates. **(B)** Representative images of adult plants of each transgenic line and the WT. **(C)** Confocal laser scanning microscopy images of the RS40-GFP and RS41-GFP proteins from transgenic plants grown in the dark for 3 days.

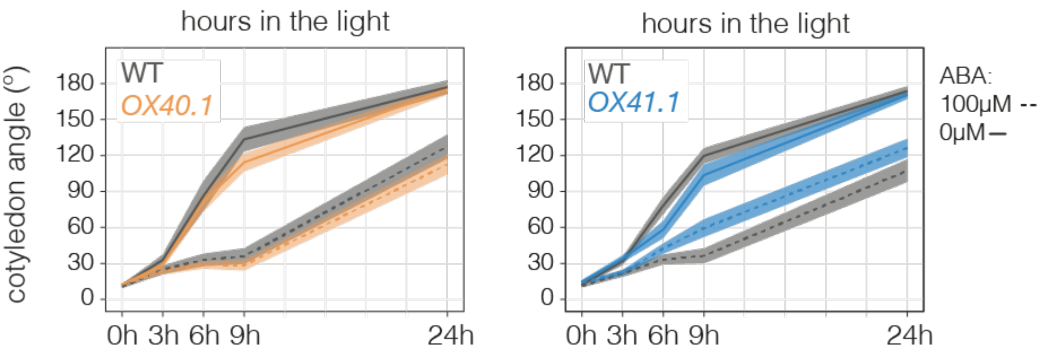


**Appendix Fig. S13. Light and ABA regulation of cotyledon opening in plants overexpressing *RS40* or *RS41*.** Quantification of cotyledon opening in the wild-type (WT) and *RS40* or *RS41* transgenic seedlings grown for 3 days in the dark and then exposed to white light for 3, 6, 9 or 24 hours (h) in the absence or presence of ABA. Thick lines and shaded areas represent respectively the median and the interquartile range of at least 70 seedlings. The experiment was repeated at least twice with similar results.

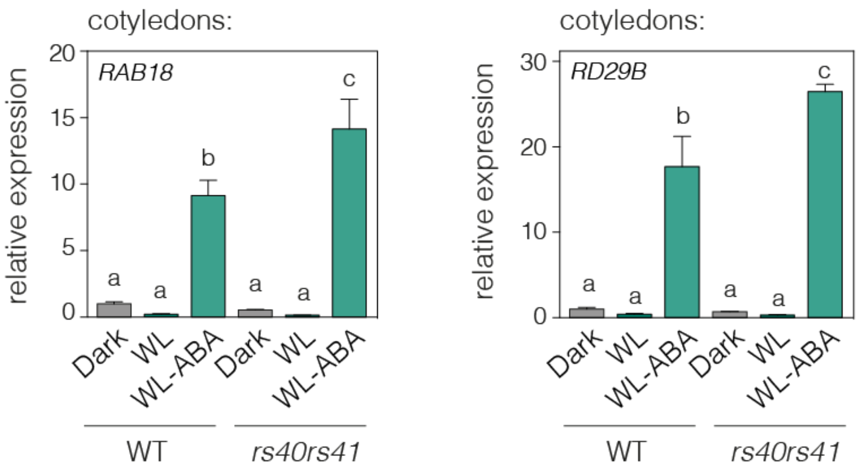


**Appendix Fig. S14. *RAB18* and *RD29B* expression levels in ABA-treated *rs40 rs41* double mutants.** RT-qPCR analysis of *RAB18* (left) and *RD29B* (right) transcript levels in wild-type (WT; Col-3) and *rs40rs41.1* cotyledons of seedlings grown for 3 days in the dark and then exposed to continuous white light (WL) for 8 hours in the absence or presence of ABA (100 µM). *PP2A* was used as a reference gene, and expression levels in the WT were set to 1. Data are the means ± SE of biological triplicates, and different letters indicate statistically significant differences between conditions by Tukey's multiple comparison test ($P < 0.05$).

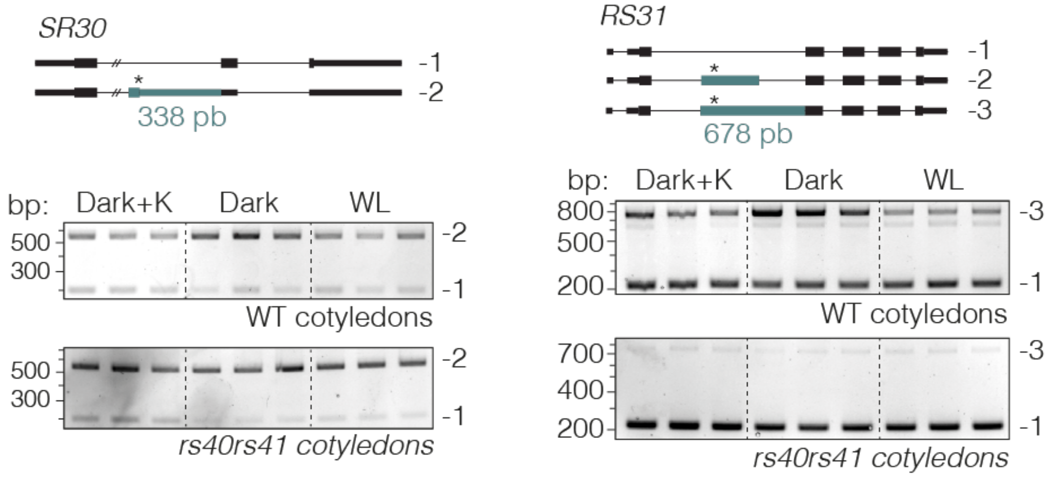


**Appendix Fig. S15. K252a regulation of *SR30* and *RS31* light- and ABA-regulated AS events.** RT-PCR analysis of *SR30* (left) and *RS31* (right) alternative transcripts in wild-type (WT; Col-3) and *rs40rs41.1* cotyledons of seedlings grown for 3 days in the dark and then exposed to continuous white light (WL) or the kinase inhibitor K252a (K; 1 μM) for 8 hours (h) in the dark. The three gel lanes for each condition are biological triplicates, and the gene diagrams on the top show the alternative sequences in green, with asterisks indicating the location of in-frame premature stop codons. bp, base pairs.

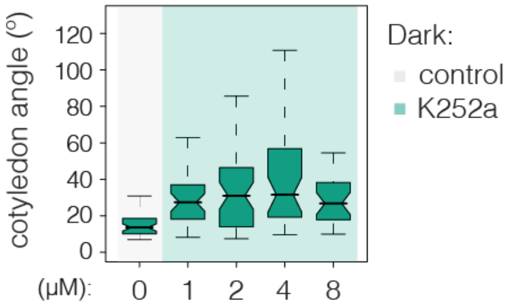


**Appendix Fig. S16. K252a regulation of cotyledon opening in etiolated seedlings.** Quantification of cotyledon opening in wild-type seedlings grown for 3 days in the dark and then treated with different concentrations of the kinase inhibitor K252a for 9 hours. Data represent the median of at least 30 seedlings.

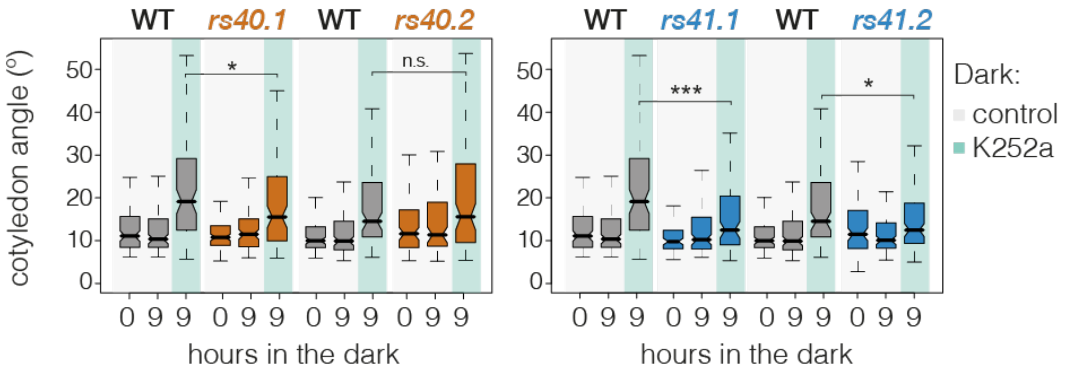


**Appendix Fig. S17. K252a regulation of cotyledon opening in the *rs40* and *rs41* single mutants.** Boxplot representation of cotyledon opening in the *rs40* (left) and *rs41* (right) single mutants and their respective wild-type (WT) seedlings (Col-0 or Col-3; see Methods for details) grown for 3 days in the dark and then treated or not with the kinase inhibitor K252a (1 μM) for 9 hours. Data represent the median of at least 80 seedlings, and asterisks indicate statistically significant differences between K252a-treated mutants and their respective WTs (Mann–Whitney test; *, $P$ < 0.05; **, $P$ < 0.01; ***, $P$ < 0.001; n.s., non-significant). The experiment was repeated at least twice with similar results.

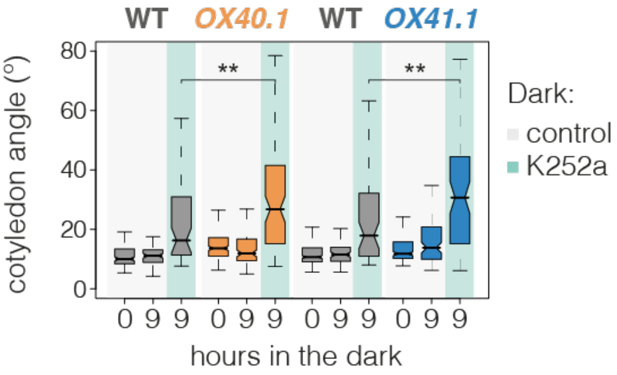


**Appendix Fig. S18. K252a regulation of cotyledon opening in plants overexpressing *RS40* or *RS41*.** Quantification of cotyledon opening in wild-type (WT) and *RS40*- or *RS41*-overexpressing seedlings grown for 3 days in the dark and then treated or not with the kinase inhibitor K252a (1 μM) for 9 hours. Data represent the median of at least 50 seedlings, and asterisks indicate statistically significant differences between treated transgenic plants and their respective WTs (Mann–Whitney test; *, $P < 0.05$; **, $P < 0.01$; ***, $P < 0.001$; ns, non-significant). The experiment was repeated at least twice with similar results.
